# Somatostatin, an *In Vivo* Binder to Aβ Oligomers, Binds to βPFO_Aβ(1-42)_ Tetramer

**DOI:** 10.1101/2020.05.11.088153

**Authors:** Eduard Puig, James Tolchard, Antoni Riera, Natàlia Carulla

**Author notes:** Corresponding authors: Eduard Puig, Natàlia Carulla.

## Abstract

Somatostatin (SST14) is strongly related to Alzheimer’s disease (AD), as its levels decline during aging, it regulates the proteolytic degradation of the amyloid beta peptide (Aβ), and it binds to Aβ oligomers *in vivo*. Recently, the 3D structure of a membrane-associated β-sheet pore forming tetramer (βPFO_Aβ(1-42)_ tetramer) has been reported. Here we show that SST14 binds selectively to the βPFO_Aβ(1-42)_ tetramer without binding to monomeric Aβ(1-42). Specific NMR chemical shift perturbations, observed during titration of SST14, define a binding site in the βPFO_Aβ(1-42)_ tetramer and are in agreement with a 2:1 stoichiometry determined by both native MS and ITC. These results enabled us to perform driven docking and model the binding mode for the interaction. The present study provides additional evidence on the relation between SST14 and the amyloid cascade, as well as positions the βPFO_Aβ(1-42)_ tetramer as a relevant aggregation form of Aβ and as a potential target for AD.

## Introduction

Increased levels of amyloid beta peptide (Aβ) and deposition of amyloid fibrils in neuronal cells constitute a critical part of the etiopathogenesis in Alzheimer’s disease (AD) (*Hardy and Higgins, 1992; Selkoe and Hardy, 2016)*. Aβ originates from the sequential cleavage of the amyloid precursor protein (APP) by the β-secretase in the extracellular space and the *γ*-secretase in the transmembrane domain (*Kimberly et al., 2003*). In solution, this hydrophobic peptide aggregates in a nucleation-dependent manner into soluble oligomers (*Haass and Selkoe, 2007*) that gradually increase in molecular-weight until insoluble fibrils are formed (*Colvin et al., 2016; Gremer et al., 2017; Wälti et al., 2016; Xiao et al., 2015*). The presence of amyloid fibrils in the extracellular space has inevitably drawn research interest in Aβ peptides to this location. However, the fact that Aβ’s origin lies within APP, a transmembrane protein (*Barrett et al., 2012*), together with numerous reported work of Aβ interacting with the cellular membrane (*Arispe et al., 1993; Bode et al., 2017; 2019; Butterfield and Lashuel, 2010; Hirakura et al., 1999; Kayed et al., 2004*), strongly suggests this environment as an alternative location for Aβ accumulation and aggregation.

We have previously studied the aggregation of Aβ within detergent micelles to mimic the membrane environment and reported the preparation (*Serra-Batiste et al., 2016*) and the three-dimensional (3D) structure (*Ciudad et al., 2019*) of a membrane-associated β-sheet pore forming tetramer (βPFO_Aβ(1-42)_ tetramer). Interestingly, the formation of this oligomer was specific for Aβ(1-42), the variant most related to AD but not Aβ(1-40) which is the variant most abundant in the brain. βPFO_Aβ(1-42)_ tetramer comprises a β-sheet core formed by six β-strands. Molecular dynamics showed that when βPFO_Aβ(1-42)_ tetramer incorporated into lipid bilayers and water molecules were able to permeate the membrane through the hydrophilic edges of the β-sheet core tetramer. This work not only represented the resolution of the first 3D structure of an Aβ oligomer but also the definition of a new mechanism of membrane disruption that could explain the neurotoxic activity of Aβ oligomers in the context of AD.

The screening of potential interactors constitutes an essential part to better understand the function of Aβ and its implication in AD. Schmitt-Ulms *et al*. recently performed an extensive screening of proteins that bound to Aβ oligomers in human brain extracts (*Wang et al., 2017*). From over 50 proteins detected, somatostatin (SST14) stood out for delaying Aβ aggregation and binding specifically to Aβ oligomers. To the best of our knowledge, the aforementioned work represents the largest Aβ monomeric and oligomeric *in vivo* interactome performed so far. The authors suggested that further investigations should be performed to improve the understanding of the SST14-Aβ interaction. In the present work, we used well established biophysical techniques to assess whether the specific Aβ oligomer binder, somatostatin-14 (SST14) bound to the βPFO_Aβ(1-42)_ tetramer.

SST14 is a cyclic tetradecapeptide that is produced in neuroendocrine cells in the hypothalamus as well as in other tissues, including pancreas, intestinal tract and regions of the central nervous system (*Morisset, 2017; Reichlin, 1983*). In a clinical context, SST14 is the neuropeptide that exhibits the strongest depletion in both the brain and cerebrospinal fluid (CSF) of AD patients (*Davies et al., 1980; Hayashi et al., 2011*). The relation to AD was further described by the work of Saido *et al.* as they found that SST14 regulates the metabolism of Aβ in the brain through the modulation of neprilysin which catalyses its proteolytic degradation (*Saito et al., 2005*). Moreover, a positive correlation between SST14 and Aβ(1-42) levels was established in the CSF of elderly patients with mild cognitive impairment (*Duron et al., 2018*). Undoubtedly, previous work in the literature has established a strong link between SST14, Aβ and AD. Therefore, studying the potential interaction between SST14 and βPFO_Aβ(1-42)_ tetramer could deliver evidence to better understand the role of SST14 in the context of AD and point towards the relevance of βPFO_Aβ(1-42)_ tetramer in a biological context.

## Results

### SST14 coelutes with βPFO_Aβ(1-42)_ tetramers

We initially relied on size exclusion chromatography (SEC) (*Bai, 2015; Bianchi et al., 2018; Rasmussen et al., 2011*) to characterize the potential interaction between SST14 and βPFO_Aβ(1-42)_ tetramer in a membrane mimicking environment. As control samples, we followed the evolution of βPFO_Aβ(1-42)_ tetramer formation and SST14 independently, following its incubation in the dodecyl phosphocholine (DPC) solution used as a membrane mimicking environment (Supporting information; Figure S1A, B). Analysis of both samples showed that βPFO_Aβ(1-42)_ tetramers and SST14 eluted, respectively, at 13.5 mL and 17.5 mL. In the case of the SST14 sample, its evolution over time revealed the appearance of wide peaks near the void volume, which were attributed to aggregated forms as previously reported for this peptide (*Anoop et al., 2014*). Coincubation of Aβ(1-42) with SST14 under conditions that lead to βPFO_Aβ(1-42)_ tetramer formation resulted in an increase of 65% of the area under the peak assigned to βPFO_Aβ(1-42)_ tetramers (Figure 1A). Such a change could be explained either due to an increase in the formation of βPFO_Aβ(1-42)_ tetramer or the binding of SST14. Interestingly, we did not observe precipitates when both peptides were coincubated suggesting that the interaction between them increased the stability of SST14 in a membrane-mimicking environment. To assess whether binding occurred specifically during βPFO_Aβ(1-42)_ tetramer formation, we first incubated Aβ(1-42) alone for 24 h under conditions that lead to βPFO_Aβ(1-42)_ tetramer formation and then added SST14. Analysis of this sample by SEC resulted in a 20% increase of the area under the peak assigned to βPFO_Aβ(1-42)_ tetramer (Supporting information, Figure S1D). This result suggested that SST14 binding was not exclusively occurring during βPFO_Aβ(1-42)_ tetramer formation but also when putting in contact the two binding partners after the oligomer was assembled.

**Figure 1.**
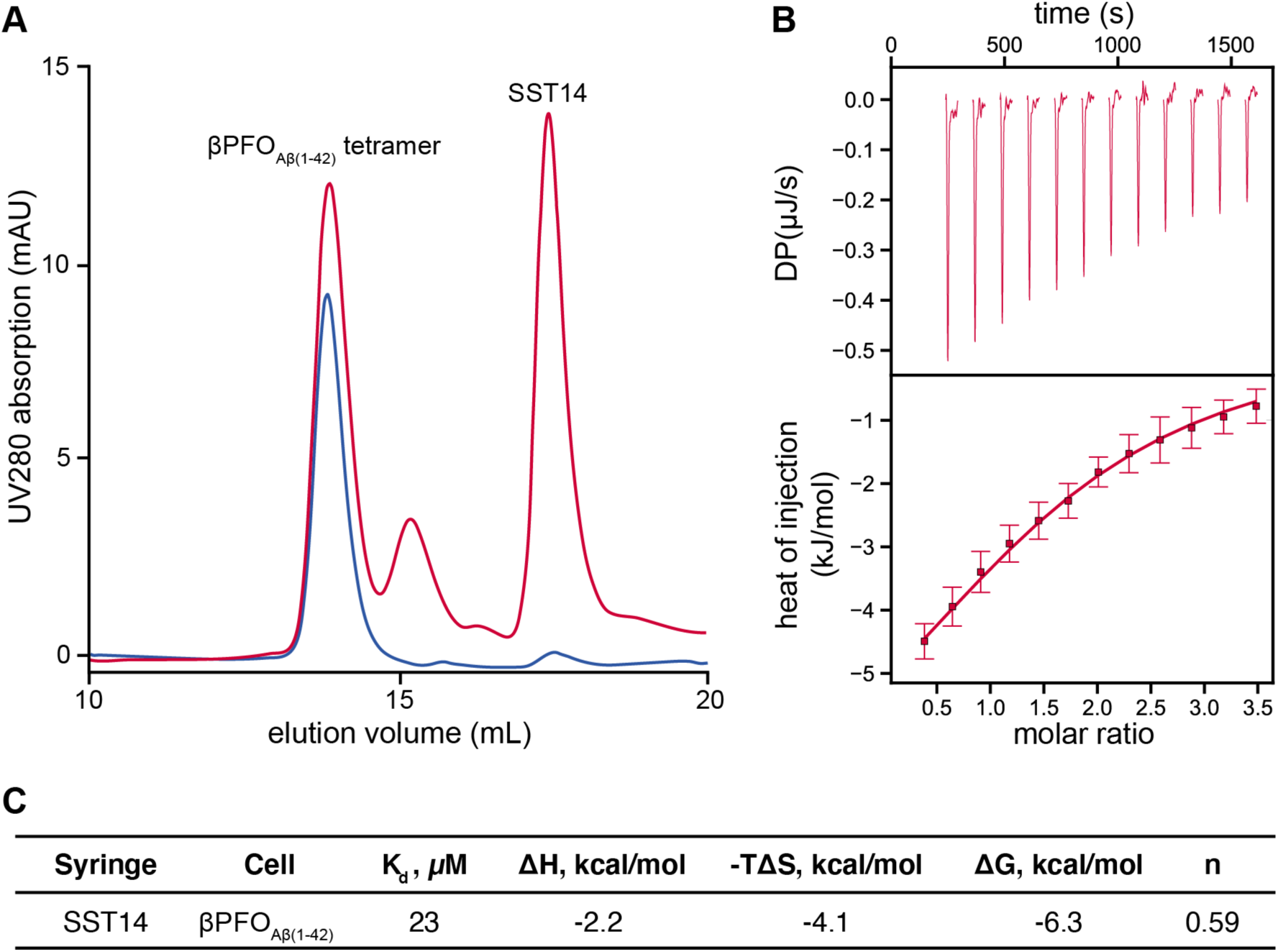
SST14 binding to βPFO_Aβ(1-42)_ tetramer assessed by SEC and ITC. (**A**) SEC elution profile for βPFO_Aβ(1-42)_ tetramer after 24 hours of its formation in the absence (blue) and in the presence of SST14 (red). The peaks have been labeled with the elution volume of βPFO_Aβ(1-42)_ tetramer and SST14. (**B**) ITC thermogram (top) and analysis of the fitted binding isotherm (bottom) for βPFO_Aβ(1-42)_ tetramer titrated with SST14. (**C**) Thermodynamic binding parameters of the interaction determined from ITC experiments at 25°C and pH 9.

### SST14 interacts with the Aβ(1-42) tetramer at a 2:1 ratio

To further study the interaction between βPFO_Aβ(1-42)_ tetramer and SST14, we used isothermal titration calorimetry (ITC) to measure the heat exchange and obtain information about the energetic profile of the binding event. Titration of SST14 to the βPFO_Aβ(1-42)_ tetramer showed an exothermic interaction with a K_D_ of 23 µM and a stoichiometry of approximately 2:1 (Figure 1B, C). Such a binding ratio would indeed be in agreement with the symmetric structure of the βPFO_Aβ(1-42)_ tetramer. However, we could not exclude the possibility of SST14 interacting with remaining monomeric Aβ(1-42) in the sample.

To better understand the specificity and stoichiometry of the interaction we analyzed the sample using native mass spectrometry (MS). This technique uses non-denaturing conditions to prepare the sample and soft ionization methods (such as electrospray ionization (ESI)) to preserve the non-covalent interactions within (βPFO_Aβ(1-42)_ tetramer) and between (βPFO_Aβ(1-42)_ tetramer-SST14) molecular complexes (*Gupta et al., 2017; Laganowsky et al., 2013*). To prepare the βPFO_Aβ(1-42)_ tetramer sample for MS analysis, we used lauryldimethylamine *N*-oxide (LDAO) instead of DPC to mimic the membrane environment as this detergent also supports βPFO_Aβ(1-42)_ tetramer formation and is compatible with MS analysis (*Reading et al., 2015*). Direct infusion of the resulting sample using nanoESI-MS delivered a clean spectrum displaying four consecutive charge states for the tetramer (+3, +4, +5 and +6) confirming that it was the major species in the sample (Figure 2A; Supporting information, Table S2). Infusion of a βPFO_Aβ(1-42)_ tetramer sample prepared in the presence of SST14 revealed consecutive charge states that were assigned to one (+3, +4 and +5) and two (+4, +5 and +6) SST14 molecules bound to the βPFO_Aβ(1-42)_ tetramer (Figure 2B; Supporting information, Table S2). Both ITC (Figure 1C) and native MS (Figure 2B) data pointed towards a 2:1 ratio for SST14 and βPFO_Aβ(1-42)_ tetramer interaction. Moreover, we did not observe any consecutive charge states corresponding to monomeric Aβ(1-42) bound to one or two SST14 molecules in agreement with SST14 binding specifically to oligomeric forms of Aβ (Supporting information; Figure S2, Table S1).

**Figure 2.**
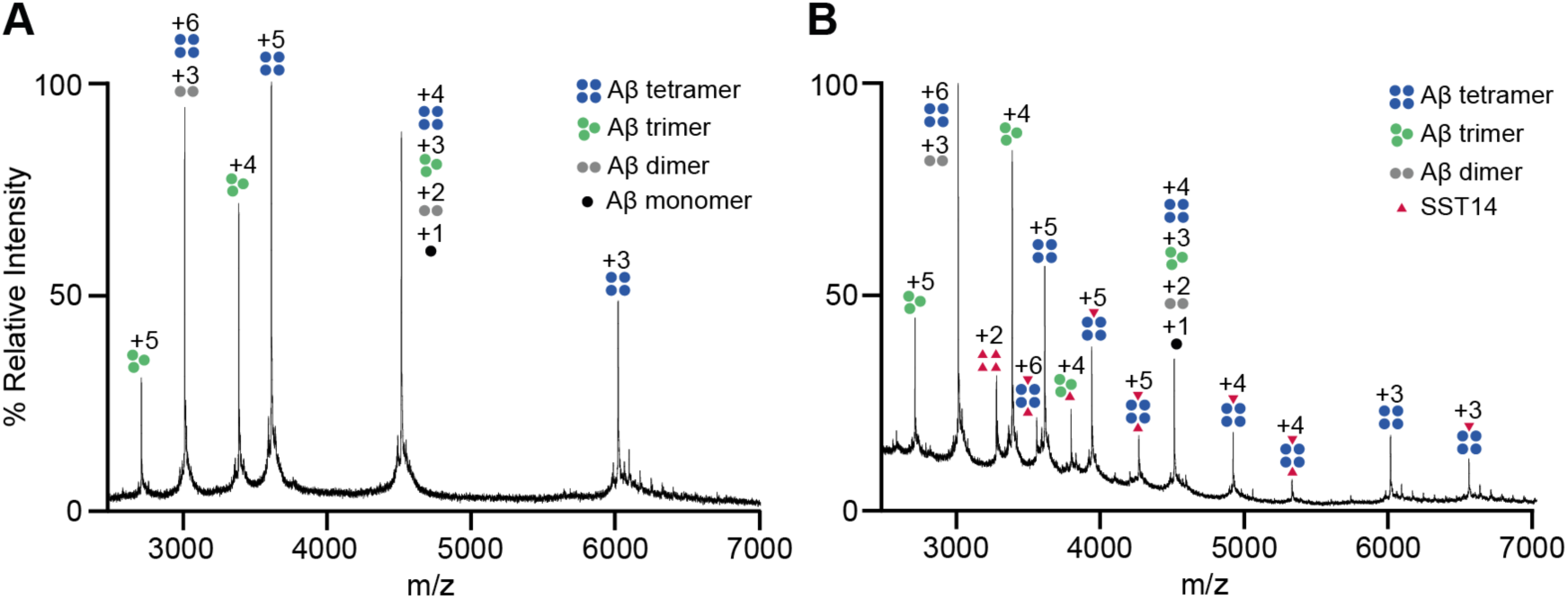
SST14 binding to βPFO_Aβ(1-42)_ tetramer assessed by native MS. (**A**) Electrospray ionization MS (ESI-MS) spectrum of βPFO_Aβ(1-42)_ tetramer (150 μM Aβ(1-42), 7.2 mM LDAO, 200 mM Ammonium Carbonate, pH 9.0 incubated for 24 hours). (**B**) ESI-MS spectrum of βPFO_Aβ(1-42)_ tetramer coincubated with SST14 (150 μM Aβ42, 150 μM SST14, 7.2 mM LDAO, 200 mM Ammonium Carbonate, pH 9.0 incubated for 24 hours). Charge states corresponding to SST14; Aβ(1-42) monomer, dimer, trimer, and tetramer are indicated with schematic drawings and labelled, respectively, in red, black, grey, green, and blue.

### SST14 binds to the flexible edges of the Aβ(1-42) tetramer

The results obtained by SEC indicated that the binding event was stable over 24 h in the membrane-mimicking environment (Supporting information, Figure S1C), which encouraged us to further study the interaction by solution NMR. We therefore decided to pursue a deeper characterization by titrating SST14 into a ^15^N-βPFO_Aβ(1-42)_ tetramer sample and perform 2D [^1^H,^15^N]-SOFAST-HMQC experiments over time. The βPFO_Aβ(1-42)_ tetramer consists of a six-stranded β-sheet comprising two types of Aβ(1-42) subunits referred to orange and green, respectively (Supporting information, Figure S3). The orange subunit contributes with two β-strands (β1 and β2) and the green subunit contributes with one β-strand (β3) and a small α-helix (α1) (*Ciudad et al., 2019*).

The spectrum for βPFO_Aβ(1-42)_ tetramer displayed a well-dispersed set of signals as previously reported for this sample (Figure 3A) (*Serra-Batiste et al., 2016*). Upon addition of SST14 to the NMR sample, several changes in chemical shifts were observed in the resulting spectrum indicating that SST14 bound to βPFO_Aβ(1-42)_ tetramer. Indeed, specific shift changes in residues V12, F20, V24 G29, V40 and A42 of the orange subunit and in residues V12, V18, A21, E22, D23, G29, I41 and A42 of the green subunit of the βPFO_Aβ(1-42)_ tetramer were observed (Figure 4A; Supporting information, Figure S4). Chemical Shift Perturbations (CSPs) were considered significant if the values were greater than the standard deviation (*σ*) of the Euclidean chemical shift change represented as a grey dashed line (Figure 4A) (*Williamson, 2013*). Moreover, observation of the smooth migration of A42 (folded peak) and G29 from the free position in the spectrum (blue) to the bound position (red) indicated that the exchange rate of binding occurred in the fast regime (Figure 3B) (*Williamson, 2013*).

**Figure 3.**
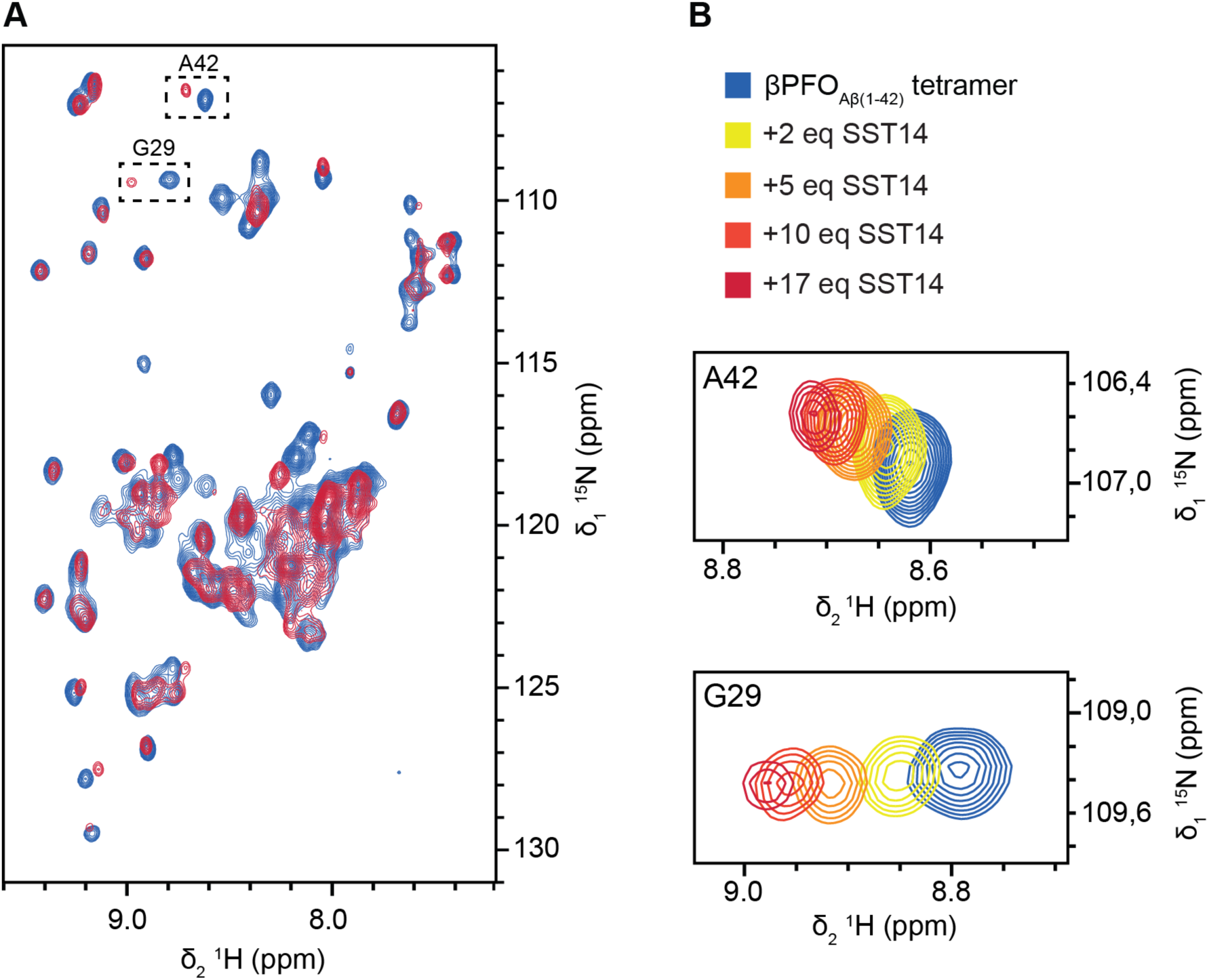
NMR titration of SST14 to βPFO_Aβ(1-42)_ tetramer. (**A**) Two-dimensional [^15^N, ^1^H]-SOFAST-HMQC spectra of βPFO_Aβ(1-42)_ tetramer (230 μM Aβ(1-42), 7.71 mM d_38_-DPC, 10 mM d_12_-Tris·DCl, pH 8.5) alone (blue) and in the presence of 17 equivalents of SST14 (red). (**B**) Close-up views of selected residues A42 (folded peak) and G29 from the titration in the presence of 0 (blue), 2 (yellow), 5 (orange), 10 (coral) and 17 (red) equivalents (eq.) of SST14.

**Figure 4.**
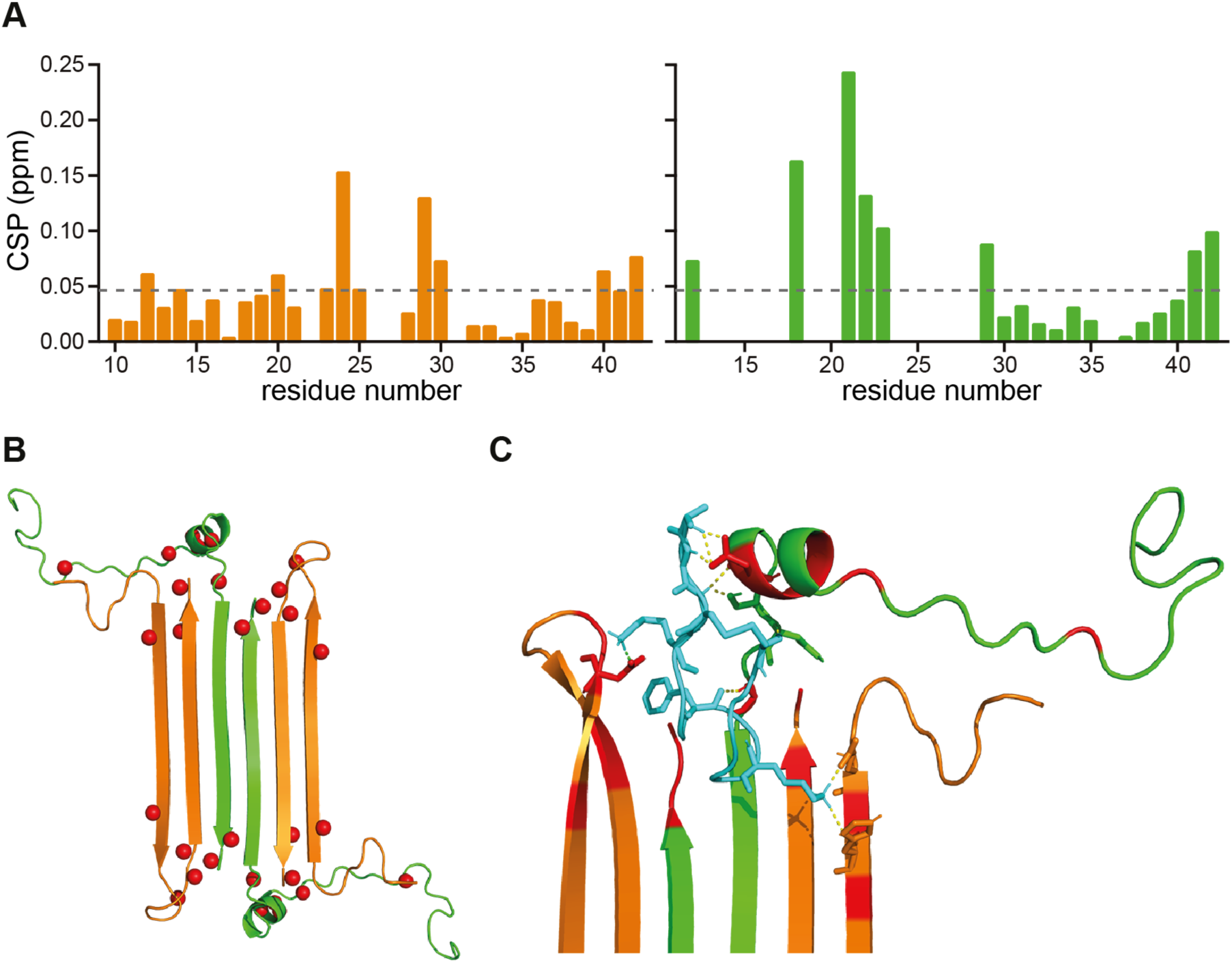
Binding site of SST14 to βPFO_Aβ(1-42)_ tetramer. (**A**) Representation of the CSP of the residues within the orange and green subunits of the βPFO_Aβ(1-42)_ tetramer induced by the presence of SST14. The grey dashed line indicates the threshold dictated by the standard deviation (σ). (**B**) Representation of the residues affected by chemical shift changes (red spheres) in the presence of SST14 within the 3D structure of the βPFO_Aβ(1-42)_ tetramer (PDB code 6RHY). (**C**) Best-ranked structure proposing a binding mode of SST14 (cyan, PDB code 2MI1) with βPFO_Aβ(1-42)_ tetramer (orange and green). Residues introduced as ambiguous interaction restraints (AIRs) are colored in red and hydrogen bonds involved in the binding are represented as yellow dashed lines.

The CSPs induced by SST14 were represented in the βPFO_Aβ(1-42)_ tetramer 3D structure (PDB code 6RHY) where the amide protons of the affected residues are represented as red spheres (Figure 4B). The affected residues showed to be close in space and defined a specific binding site within the tetramer structure. These CSPs were used to perform driven docking of SST14 with the βPFO_Aβ(1-42)_ tetramer structure using the high ambiguity driven docking approach (HADDOCK) (*Dominguez et al., 2003*). The best-ranked structure that we obtained suggested a binding mode of SST14 in the βPFO_Aβ(1-42)_ tetramer where the peptide interacted with the flexible edges of the tetramer and interestingly also tightly with the alpha helix of the green subunit (Figure 4C; Supporting information, Figure S5).

The binding site defined in our study, enabled us to rationalize the specificity of the interaction between both entities since the disposition of the residues in space for the βPFO_Aβ(1-42)_ tetramer is completely different than that for monomeric Aβ(1-42). The localization of the site was also in accordance with the 2:1 stoichiometry of the binding observed by ITC and native MS. Indeed, the symmetric topology of the tetramer contains two possible binding sites in the superior and inferior flexible ends that are solvent-exposed and thus, accessible to SST14.

## Discussion

In summary, our findings show that SST14 binds βPFO_Aβ(1-42)_ tetramers with an affinity in the low micromolar range. Native MS experiments prove the binding to be specific for this oligomeric form with a 2:1 SST14: βPFO_Aβ(1-42)_ tetramer stoichiometry, in accordance with ITC data.. Our NMR experiments reveal two symmetric binding sites near the flexible ends of the tetramer. Restraint-driven *in silico* docking enables us to propose a binding mode of SST14 to the tetramer structure. Altogether, we conclude that SST14, an *in vivo* binder to Aβ oligomers (*Wang et al., 2017*), specifically binds to βPFO_Aβ(1-42)_ tetramer.

We observed an important difference when comparing our results with previously reported work on the binding of SST14 to soluble oligomers of Aβ(1-42) (*Wang et al., 2017*). Indeed, work by Schmitt-Ulms and collaborators postulated that SST14 did not bind to Aβ(17-42) oligomers which led them to conclude that the N-terminus was involved in the binding site. Our data, on the contrary, suggests that residues 18-29 are mainly involved in the binding with special emphasis on the ones forming the short alpha helix, residues L17 to F20. We recently showed that while Aβ(1-42) incorporates both as the orange and green subunit in the βPFO_Aβ(1-42)_ tetramer arrangement, Aβ(17-42) only incorporates as the green subunit (*Ciudad et al., 2019*), which prevents Aβ(17-42) to form βPFO_Aβ(1-42)_ tetramer by itself. Thus, for βPFO_Aβ(1-42)_ tetramer to form it is required that at least 50% of the peptides contain the N-terminus. These results evidence that using shortened versions of proteins can have a huge impact in protein self-assembly and structure. Moreover, work by Schmitt-Ulms *et al.* was performed on soluble oligomers while ours on a membrane-associated oligomer. Therefore, the binding to SST14 may be different for each oligomer type. The authors also emphasized the importance of W8 of SST14 for the binding to occur. Indeed, this residue has been described to play an important role in the activity of the peptide when binding SSTRs (*Veber et al., 1978*). Our proposed binding mode does not involve direct interactions of βPFO_Aβ(1-42)_ tetramer with W8 of SST14, although it is close in space to V40 and A42 (Figure 4A and Supporting information, Figure S5C) suggesting that W8 of SST14 could also be involved in the interaction of SST14 with the membrane-associated Aβ(1-42) tetramer.

In the AD context, the critical role of SST14 in the metabolism of Aβ(1-42) through the regulation of neprilysin (*Saito et al., 2005*) inevitably points towards the potential degradation of oligomeric forms of Aβ(1-42). Two important conclusions of the work by Saido and collaborators are the location of the SST14-neprylisin interaction situated near or in the cellular membrane and the fact that somatostatin-regulated neprilysin activity selectively depleted Aβ(1-42) but not Aβ(1-40). Interestingly, the aforementioned facts also apply to βPFO_Aβ(1-42)_ tetramer since it is able to incorporate into membranes (*Ciudad et al., 2019*) and is exclusively formed by Aβ(1-42) but not Aβ(1-40) (*Serra-Batiste et al., 2016*). Proteolytic activities are tightly controlled biological processes that can be regulated at different levels such as through the formation of an activation complex *(Turk, 2006*). Therefore, we cannot exclude that binding of SST14 to the βPFO_Aβ(1-42)_ tetramer could induce its degradation.

In the present study, we show at a structural level how SST14, which has been reported to bind to Aβ oligomers in human brain extracts, also binds to βPFO_Aβ(1-42)_ tetramer. We think these results strengthen the relation of SST14 with the amyloid cascade and due to the clear implication of SST14 in AD, it positions βPFO_Aβ(1-42)_ tetramer as relevant oligomer form of Aβ and as a potential target for AD.

## Acknowledgement

This study was supported by Spanish Ministry of Economy, Industry and Competitiveness (MINECO, SAF2015-68789 to N.C. and CTQ2017-87840-P to A.R.), the Fondation Recherche Médicale (AJE20151234751) and the Counseil Régional d’Aquitaine Limousin Poitou-Charentes (1R30117-00007559). E.P. was a Ph.D. fellow funded by MINECO (FPI). We thank institutional funding from MINECO through the Centres of Excellence Severo Ochoa Award, and IRB Barcelona through the CERCA Programme of the Catalan Government. We acknowledge Mass Spectrometry and Proteomics Core Facility of IRB Barcelona, the NMR facility at IECB and Cameron Mackereth for his help with the ITC experiments and analysis.

