## supplementary_info for "Somatostatin, an *In Vivo* Binder to Aβ Oligomers, Binds to βPFO_Aβ(1-42)_ Tetramer"

### Somatostatin, an *In Vivo* Binder to Aβ Oligomers, Binds to $\beta PFO_{A\beta(1-42)}$ Tetramer

[3]Departament de Química Inorgànica i Orgànica. Universitat de Barcelona, Martí i Franqués 1, 08028, Barcelona, Spain

[*]Corresponding authors:

Eduard Puig,

Natàlia Carulla,

### Supporting Information

**Materials and Methods**

**Author Contributions**

**Supporting References**

### Materials and Methods

#### Reagents

DPC and LDAO were purchased, respectively, from Cube Biotech and Anatrace. Deuterated detergents were purchased from Cambridge Isotope Laboratories. Culture media, antibiotics and vitamins were purchased from Duchefa Biochemie. All other reagents were supplied by Sigma-Aldrich unless otherwise stated.

#### Synthetic SST14 peptide

SST14 was synthesized using solid phase peptide synthesis (SPPS) by BCN Peptides, S.A. (www.bcnpeptides.com) (Barcelona, Spain).

#### Synthetic Aβ(1-42) peptide

Aβ(1-42) was synthesized by Dr. James I. Elliott (New Haven, CT, USA) using SPPS.

#### Recombinant expression of $^{15}N$ Aβ(1-42) peptide

The recombinant expression of $^{15}N$-Aβ(1-42) was done as published previously by our group using autoinduction fused to SUMO protein (***Serra-Batiste et al., 2017***). In short, Rosetta (DE3) pLysS *E. coli* cells (Novagen) were transformed with the expression vector and grown overnight at 37°C on Luria Bertani (LB)-agar plates containing 1% glucose. All cell cultures were also supplemented with 35 µg/mL chloramphenicol and 50 µg/mL kanamycin. Single colonies were picked and grown overnight in 12.5 mL LB, 1% glucose. The pre-cultures were centrifuged at 3,000 *g* for 10 min at 25°C. Each pellet was transferred to 0.5 L $^{15}N$-labeled P-5052 auto-inducing media with the appropriate antibiotics using a 3 L Erlenmeyer flask. The resulting cultures were grown for 6 h at 37°C and 180 rpm. The temperature was then lowered to 25°C, and the culture was incubated 22 h more at 180 rpm. The cells were then harvested by centrifugation at 9,000 *g* for 15 min at 4°C and then frozen at -80°C.

#### Purification of recombinant $^{15}N$ Aβ(1-42) peptide into monomeric form

The cell pellet was resuspended with 6 mL buffer A (300 mM NaCl, 50 mM $Na_3PO_4$, 20 mM imidazole, 1% Tween-20, and 1 mM Tris(2-carboxyethyl)phosphine (TCEP) at pH 8.0), supplemented with half a pill of ethylenediaminetetraacetic acid (EDTA) free Complete protease inhibitor (Roche) and 1 mg DNAse (Roche) per gram of cells. The resuspended cells were lysed using a cell disruptor (Constant Systems Ltd. U.K.) operating at 20,000 psi. The cell extract was then centrifuged at 30,000 *g* for 30 min at 4°C and the resulting supernatant was filtered using a 0.45 µm before Immobilized Metal Affinity Chromatography (IMAC).

The supernatant was loaded at 1 mL/min onto a HisTrap HP 5-mL Ni column (GE Healthcare), previously equilibrated with the aforementioned buffer A. The washing was done with buffer B (300 mM NaCl, 50 mM sodium phosphate, 40 mM imidazole, 0.05% Tween-20 and 1 mM TCEP at pH 8.0) for 10-15 column volumes, until UV absorbance was stable. Elution was done following a 3-step method: (a) 15 mL linear gradient from 0 to 15% of buffer C (300 mM NaCl, 50 mM sodium phosphate, 500 mM imidazole, 0.05% Tween-20 and 1 mM TCEP pH 8.0), followed by (b) a 20 mL isocratic step at 15% buffer C and (c) a second isocratic step at 100% buffer C until UV absorbance was stable

To remove imidazole from the samples, they were buffer-exchanged using a HiPrep 26/10 desalting column (GE Healthcare) equilibrated with 50 mM ammonium carbonate and 1 mM TCEP. The concentration and purity of Aβ(1-42) was determined with reversed phase high performance liquid chromatography (RP-HPLC). The cleavage of Aβ(1-42) from the SUMO fusion tag required incubation overnight at 4°C with SUMO protease in a 1:50 [SUMO]:[Aβ(1-42)] ratio. The yield of the cleavage and the concentration of Aβ(1-42) were determined by RP-HPLC. Aliquots containing 3.75 mg of Aβ(1-42) were lyophilized before solubilization with 6.8 M guanidinium thiocyanate (Gdn·SCN) to reach 2.5 mg Aβ(1-42)/mL and sonicated for 5 min in an ice bath. Samples were then further diluted with MilliQ water down to 1.5 mg Aβ(1-42)/mL, 4 M Gdn·SCN and centrifuged at 10,000 g for 6 min at 4°C. Finally, 2.5 mL were injected into a HiLoad Superdex 30 prep grade column (GE Healthcare), previously equilibrated with 50 mM ammonium carbonate, and eluted at 4°C at a flow rate of 1 mL/min. The peaks corresponding to SUMO and monomeric Aβ(1-42) were collected separately and their purity and concentration were determined by RP-HPLC. Fractions with reduced purity (<90%) were lyophilized again and subjected to a second SEC with prior Gdn·SCN solubilization. Purified Aβ(1-42) aliquots were lyophilized and stored at -20°C until use.

#### Purification of synthetic Aβ(1-42) peptide into monomeric form

10 mg of Aβ(1-42) peptide obtained from SPPS were dissolved in 6.8 M Gdn·SCN (Life Technologies) at 8.5 mg/mL and sonicated for 5 min in a heated bath. Subsequently, the sample was diluted down to 5 mg/mL of Aβ(1-42) and 4 M Gdn·SCN with $H_2O$. It was then centrifuged at 10,000 g for 6 min at 4°C and spun with a 0.45-µm Millex filter (Millipore). The resulting Aβ(1-42) solution was injected into a HiLoad Superdex 75 prep grade column (GE Healthcare). The column was equilibrated with 50 mM ammonium carbonate at pH 9 and

eluted at 4°C at a flow rate of 1 mL/min. The peak attributed to monomeric Aβ(1-42) was collected and the concentration was determined by RP-HPLC. Purified Aβ(1-42) aliquots were lyophilized and stored at -20°C until use.

#### Quantification of Aβ(1-42) peptide

The concentration of monomeric Aβ(1-42) was determined by RP-HPLC (Waters Alliance 2695 equipped with 2998 photodiode array detector). RP-HPLC analysis was done using a Symmetry 300 C4 column (4.6 × 150 mm, 5 µm, 300 Å; Waters) at a flow rate of 1 ml/min and a linear gradient from 0 to 60 % B in 15 min (A = 0.045 % trifluoroacetic acid (TFA) in water, and B = 0.036 % TFA in acetonitrile) at 60°C. A calibration curve correlating the area under the peak and the concentration of Aβ(1-42) was generated based on a Aβ(1-42) solution previously quantified by amino acid analysis.

#### Preparation of $\beta PFO_{A\beta(1-42)}$ tetramer

$\beta PFO_{A\beta(1-42)}$ tetramer was prepared from lyophilized synthetic monomeric Aβ(1-42) samples dissolved in 5.5 mM DPC, 10mM Tris and adjusted to pH 9 reaching a final concentration of 150 µM Aβ(1-42) corresponding to a 2:1 ratio ([Aβ(1-42)]:[$DPC_m$]). Samples were incubated at 37ºC for 24 h.

#### Size Exclusion Chromatography

All the samples were filtered with a 0.45 µm spin filter (Millipore) and 10 µL were injected to a Superdex 200 increase 10/300 (GE Healthcare) column equilibrated with 3 mM DPC, 10 mM Tris·HCl, 100 mM NaCl at pH 9. Samples were eluted at 4ºC at a flow rate of 0.5 mL/min and absorbance was monitored at 214, 220, and 280 nm using an ÄKTA Pure (GE Healthcare). The following two samples were used as controls: formation of $\beta PFO_{A\beta(1-42)}$ tetramer sample prepared at 150 µM Aβ(1-42) and SST14 evolution in the same membrane mimetic environment prepared at 150 µM SST14. Coincubation of the two peptides was performed at the same concentration as the control samples, 150 µM for both Aβ(1-42) and SST14. Several SEC experiments were performed of the same samples after 0, 4, 8 and 24 h incubation (Figure S1A-C). In another case, SST14 was added to a pre-formed $\beta PFO_{A\beta(1-42)}$ tetramer sample by resuspending dry SST14 into an oligomer solution to a final concentration of 150 µM SST14 (Figure S1D, dark green trace). Chromatograms were analyzed using Unicorn (GE Healthcare) and smoothed with Illustrator CC 2019 (Adobe).

#### Isothermal Titration Calorimetry (ITC)

Experiments were performed on an $iTC_{200}$ microcalorimeter (Malvern Panalytical) at 298 K stirring at a rate of 600 rpm. The exact same batch of buffer containing 5.5 mM DPC, 10 mM Tris·HCl at pH 9 was used to resuspend the Aβ(1-42) peptide to prepare $\beta PFO_{A\beta(1\text{-}42)}$ tetramer samples and the SST14 titrant. The cell contained $\beta PFO_{A\beta(1\text{-}42)}$ tetramer at a concentration of 35 µM and the syringe contained SST14 at a concentration of 600 µM. An initial delay of 120 s was left followed by a first injection of 1 µL and subsequently 12 injections of 3 µL were titrated into the cell. The delay between each injection was 120 s. Data was processed using NITPIC to integrate the points of the titration and SEDPHAT to fit the curve (***Brautigam et al., 2016; Keller et al., 2012; Scheuermann and Brautigam, 2015; Zhao et al., 2015***). The fitted data was obtained from two independent measurements. The ITC thermogram and analysis of the fitted binding isotherm was graphed with GUSSI (***Brautigam, 2015***).

#### NanoESI-MS in non-denaturing conditions

MS samples were prepared by resuspending dry Aβ(1-42) to a final concentration of 150 *µ*M in 7.2 mM LDAO, 200 mM $(NH_4)_2CO_3$ at pH 9.0. Experiments were performed on a Synapt G1 HDMS (Waters) equipped with an Advion TriVersa NanoMate (Advion Biosciences). Positive mode of ESI was used. Typical values for source and desolvation temperature were 40°C and 300-350°C respectively. The voltages and parameters at different stages of the spectrometer were fine-tuned to optimize ion transmission as shown in table S3. Acquisitions were performed in the m/z range 1000 to 8000 with a 1.5 s scan time. External calibration was performed using singly charged ions produced by a 2 g/L solution of cesium iodide in 2-propanol/water (50/50, v/v). Spectra were smoothed (smoothing method: mean; smooth window:10; number of smooths: 2) with MassLynx V4.1 software (Waters).

#### Solution NMR

$^{15}N$ $\beta PFO_{A\beta(1\text{-}42)}$ tetramer was prepared in $d_{38}$-DPC micelles (230 µM $^{15}N$ Aβ(1-42) and 7.71 mM $d_{38}$-DPC) in 90% $H_2O$/10% $D_2O$, 10 mM $d_{12}$-Tris·DCl, pH 8.5. SST14 was titrated gradually using a 10 mM solution in the exact same buffer as used to prepare the oligomer. Concentration of SST14 at every point of the titration is detailed in table S4. All $^1H$-$^{15}N$-SOFAST-HMQC experiments were acquired at 37 °C on a 800 MHz Bruker spectrometer equipped with a cryoprobe. Data analysis was performed using CcpNMR (***Skinner et al., 2016***). To weight the relative chemical shifts of $^{15}N$ and $^1H$ nuclei, we considered a value of

0.14 for correcting factor $\alpha$ to proceed with the average Euclidean distance moved (Equation 1) (***Williamson, 2013***).

$$d = \sqrt{\frac{1}{2}[\delta_H^2 + (\alpha \cdot \delta_N^2)]} \qquad \text{Equation 1}$$

#### Docking

Somatostatin docking was carried out using the Haddock webserver (available at: https://milou.science.uu.nl/services/HADDOCK2.2/haddockserver-easy.html), using the "easy" interface (***Dominguez et al., 2003***). To conform with the input requirements, the PDB-file of the $\beta PFO_{A\beta(1\text{-}42)}$ tetramer 3D structure was used (PDB code 6RHY); no perturbations to the actual structure were made. The structure of SST14 was taken from the PDB (model 10, code: 2MI1). All $\beta PFO_{A\beta(1\text{-}42)}$ tetramer:SST14 NMR CSPs were considered as important residues during the docking calculation. However, due to the symmetry of the tetramer structure and the binary nature of the docking calculation, overall, only half of the CSPs were used – those pertaining to one half of the $\beta PFO_{A\beta(1\text{-}42)}$ tetramer, to best localize SST14 to the edge of the sheet. All 14 residues of somatostatin were considered as interaction sites.

#### Supplementary Figures

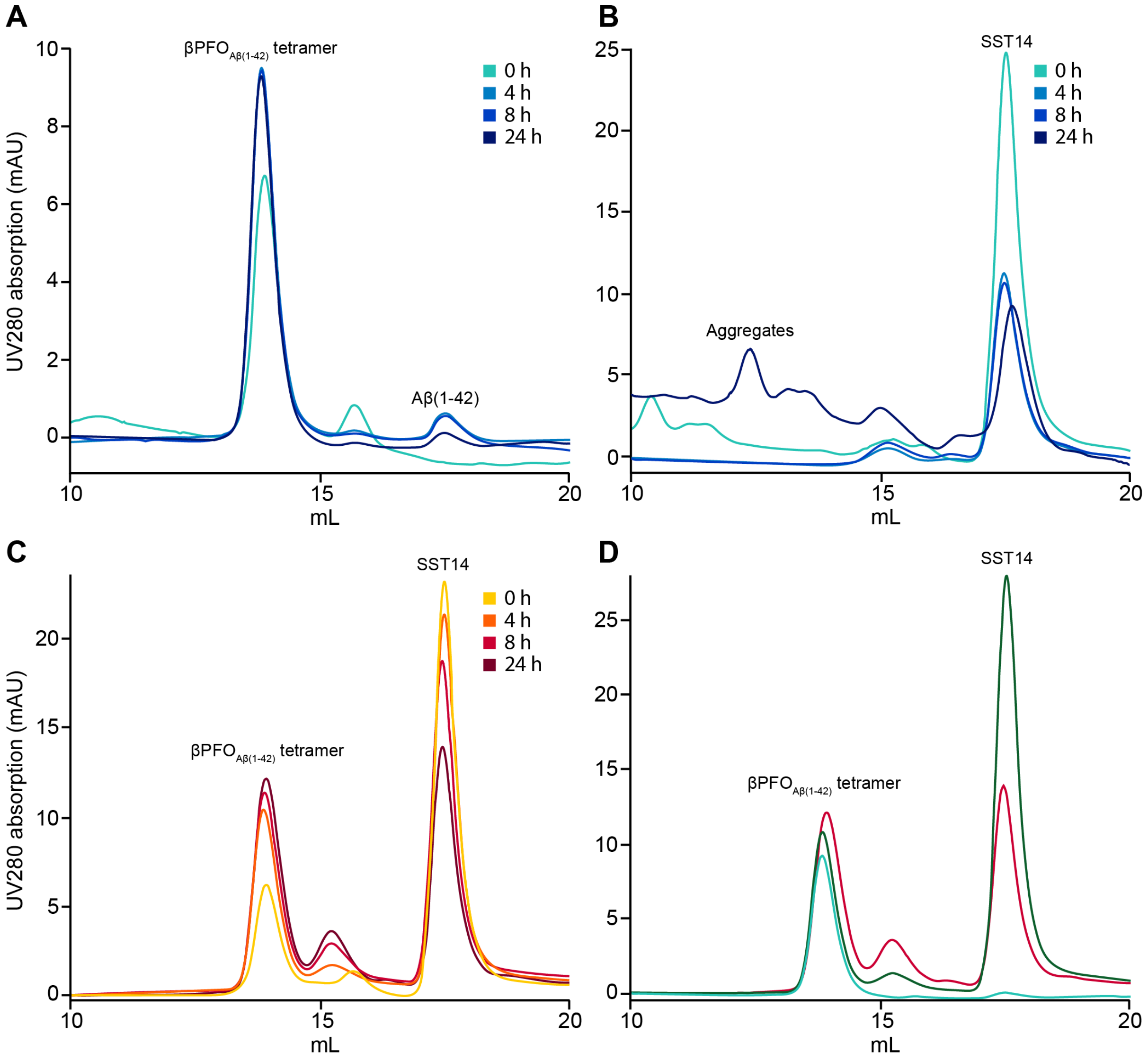

**Figure S1.** $\beta PFO_{A\beta(1-42)}$ tetramer formation in the presence and absence of SST14. (**A**) Control SEC elution profile following the $\beta PFO_{A\beta(1-42)}$ tetramer formation over 24 h in DPC buffer. (**B**) Control SEC elution profile following the evolution of SST14 over 24 h in DPC buffer. (**C**) SEC elution profile following $\beta PFO_{A\beta(1-42)}$ tetramer formation co-incubated with SST14 over 24 h in DPC buffer. (**D**) Comparison of SEC elution profiles for $\beta PFO_{A\beta(1-42)}$ tetramer after 24 h of its formation (cyan), $\beta PFO_{A\beta(1-42)}$ tetramer co-incubated with SST14 for 24 h (red) and $\beta PFO_{A\beta(1-42)}$ tetramer to which SST14 has been added after 24 h of its formation (dark green).

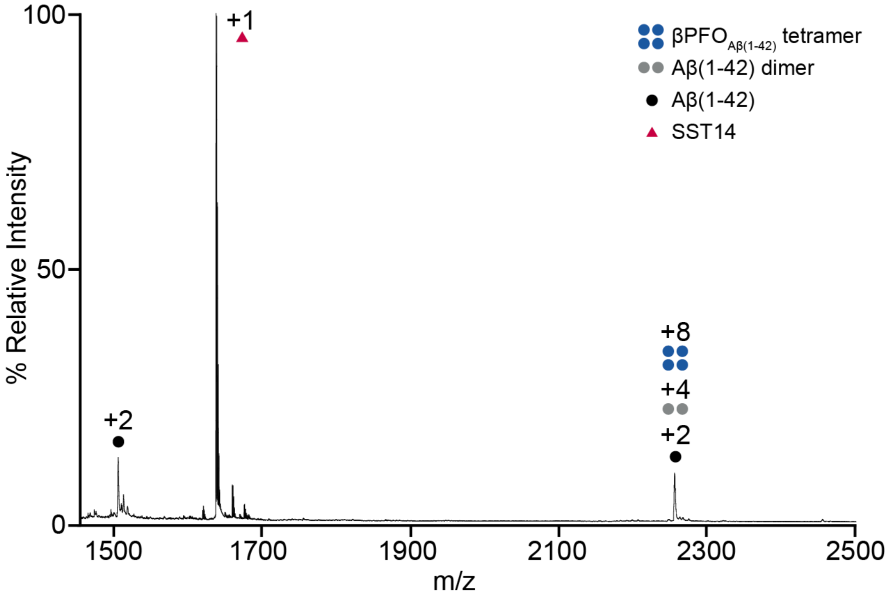

**Figure S2.** SST14 binding to Aβ(1-42) assessed by native MS. Lower mass-to-charge ESI MS spectrum of $\beta PFO_{A\beta(1-42)}$ tetramer coincubated with SST14 (150 µM Aβ42, 150 µM SST14, 7.2 mM LDAO, 200 mM Ammonium Carbonate, pH 9.0 incubated for 24 hours). Charge states corresponding to SST14; Aβ(1-42) monomer, dimer and tetramer are indicated with schematic drawings and labelled, respectively, in red, black, grey, and blue.

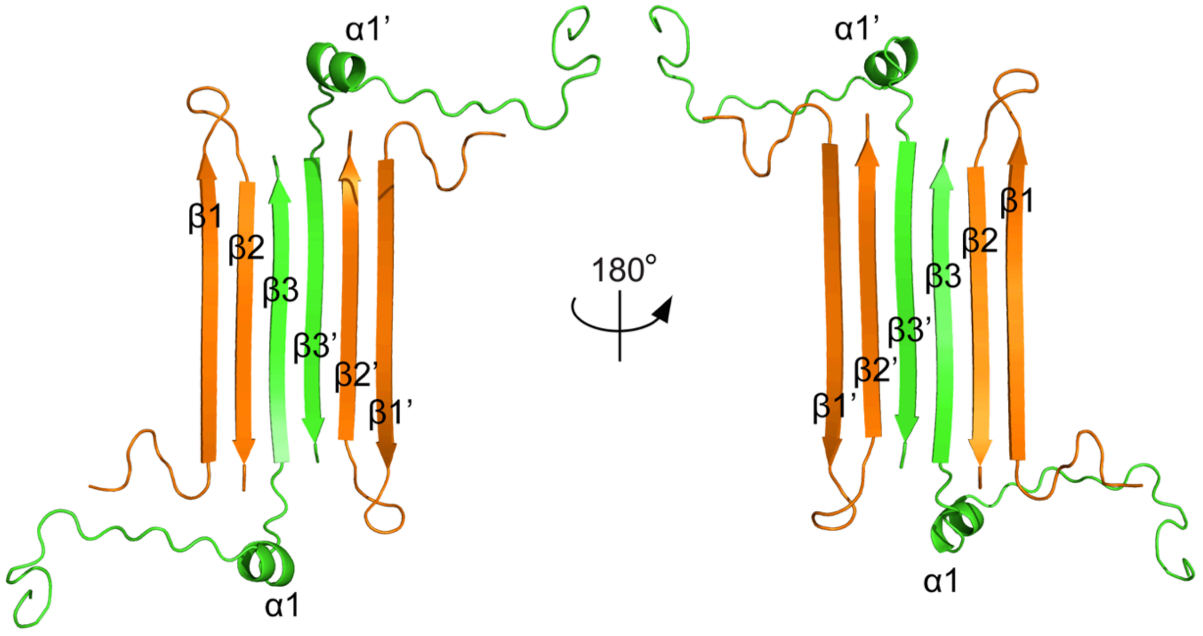

**Figure S3.** βPFO$_{Aβ(1\text{-}42)}$ tetramer structure. The βPFO$_{Aβ(1\text{-}42)}$ tetramer structure is formed by two asymmetric dimer units, which in turn are formed by two distinct Aβ(1-42) subunits, referred to as the orange and green chains. The orange subunit contributes two β-strands (β1 and β2) while the green subunit contributes with one β-strand (β3) and a short α-helix (α1).

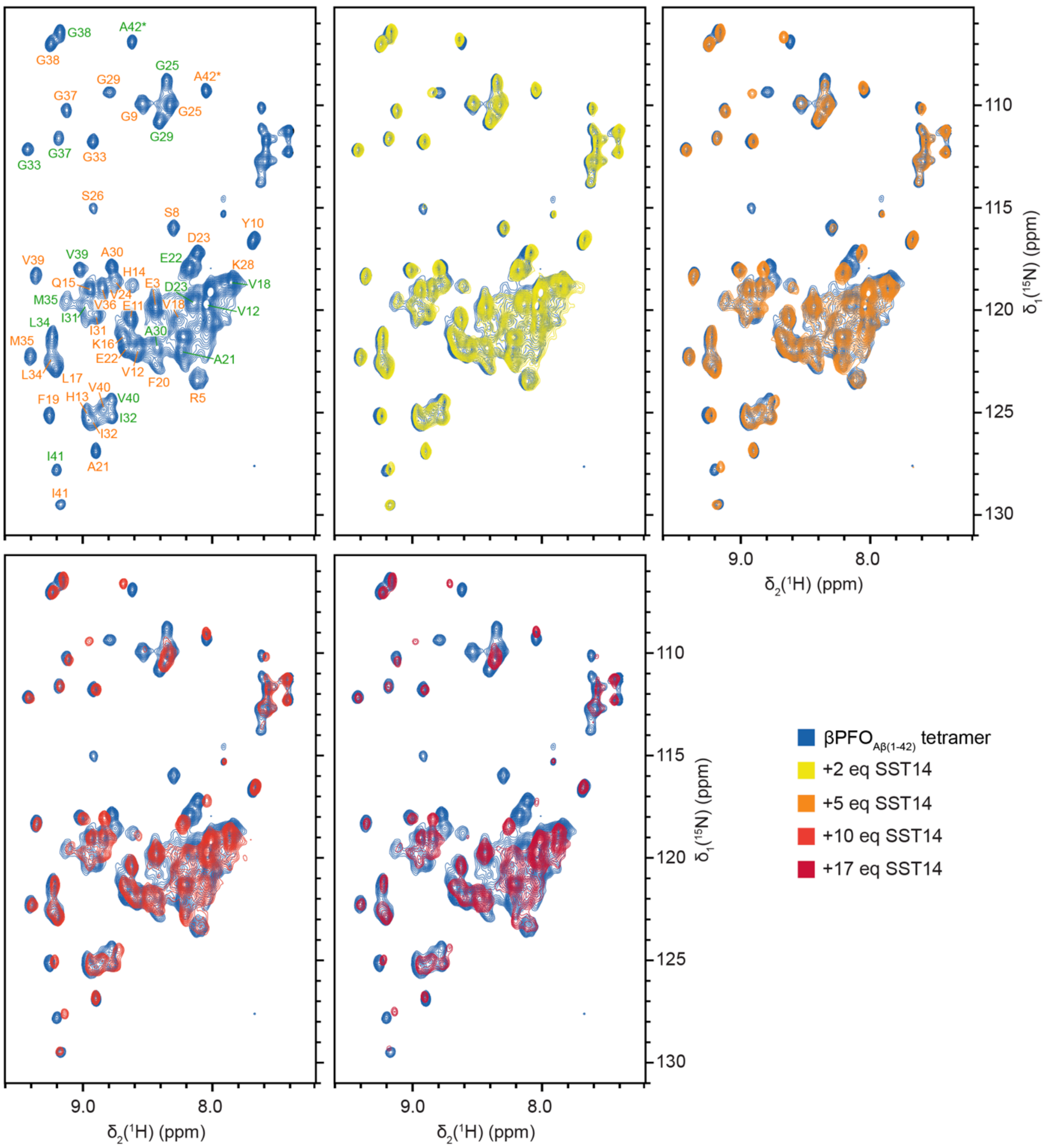

**Figure S4.** 2D [$^1$H, $^{15}$N]-SOFAST-HMQC spectrum of $\beta PFO_{A\beta(1\text{-}42)}$ tetramer in the presence of different SST14 amounts. 2D [$^1$H, $^{15}$N]-SOFAST-HMQC spectrum of $\beta PFO_{A\beta(1\text{-}42)}$ tetramer (230 µM Aβ(1-42), 7.71 mM $d_{38}$-DPC, 10 mM $d_{12}$-Tris·DCl, pH 9.0) acquired on a 800 MHz spectrometer (blue trace) where sequence-specific resonance assignments of the backbone amide groups of the two subunits are indicated in orange and green. 2D [$^1$H, $^{15}$N]-SOFAST-HMQC spectra corresponding to the titration of 2, 5, 10 and 17 equivalents of SST14 are represented in yellow, orange, coral and red traces, respectively.

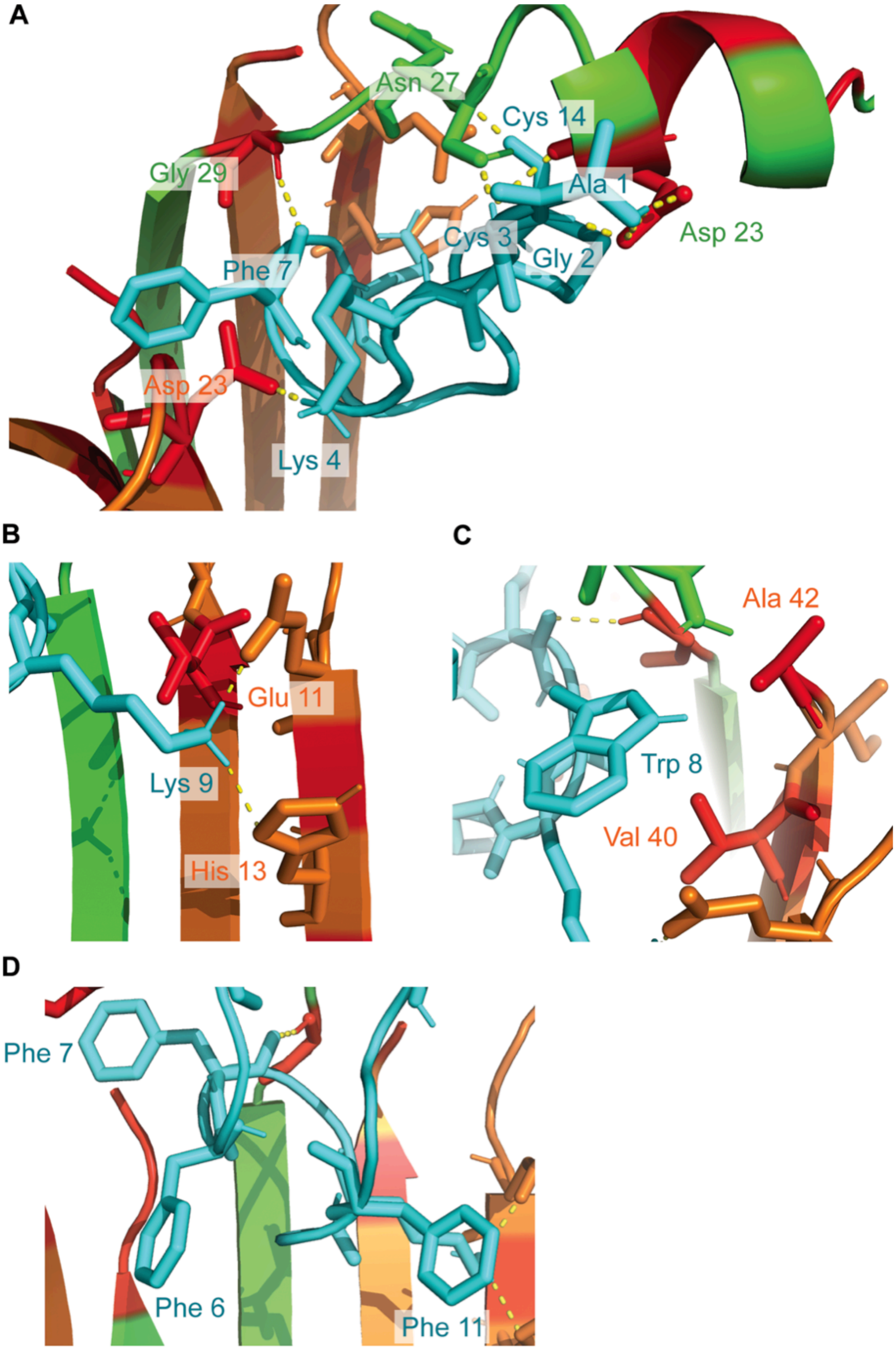

**Figure S5.** Proposed binding mode of SST14 to the βPFO$_{A\beta(1\text{-}42)}$ tetramer. (**A**) Close-up view of the proposed binding mode of SST14 (cyan) to βPFO$_{A\beta(1\text{-}42)}$ tetramer. Residues introduced as ambiguous interaction restraints (AIRs) are colored in red and hydrogen bonds involving the binding are represented as yellow dashed lines. (**B**) Interaction of Lys 9 (SST14) with Glu 11 and His 13 (tetramer, chain A). (**C**) Close-up view of Trp 8 (SST14) location. (**D**) Disposition of Phe 6, Phe 7 and Phe 11 present in SST14. The βPFO$_{A\beta(1\text{-}42)}$ tetramer structure is colored to indicate the two distinct Aβ(1-42) subunits as the orange and green chains.

#### Supporting Tables

***table S1:*** theoretically calculated m/z values for the Aβ(1-42) monomer and dimer bound to 0, 1 or 2 SST14 molecules. Signals observed in the mass spectrum were compared to these values for the assignments. Values highlighted in orange correspond to the experimentally observed charge states.

| | **Aβ(1-42) monomer** | | | **Aβ(1-42) dimer** | | |
|---|---|---|---|---|---|---|
| | 0 | 1 | 2 | 0 | 1 | 2 |
| **+1** | 4515.0 | 6152.9 | 7790.8 | 9029.1 | 10667.0 | 12304.8 |
| **+2** | 2258.0 | 3077.0 | 3895.9 | 4515.0 | 5334.0 | 6152.9 |
| **+3** | 1505.7 | 2051.6 | 2597.6 | 3010.4 | 3556.3 | 4102.3 |
| **+4** | 1129.5 | 1539.0 | 1948.4 | 2258.0 | 2667.5 | 3077.0 |
| **+5** | 903.8 | 1231.4 | 1559.0 | 1806.6 | 2134.2 | 2461.8 |
| **+6** | 753.3 | 1026.3 | 1299.3 | 1505.7 | 1778.7 | 2051.6 |
| **+7** | 645.9 | 879.8 | 1113.8 | 1290.7 | 1524.7 | 1758.7 |
| **+8** | 565.3 | 770.0 | 974.7 | 1129.5 | 1334.2 | 1539.0 |

***table S2:*** theoretically calculated m/z values for Aβ(1-42) trimer and tetramer bound to 0, 1 or 2 SST14 molecules. Signals observed in the mass spectrum were compared to these values for the assignments. Values highlighted in orange correspond to the experimentally observed charge states.

| | **Aβ(1-42) trimer** | | | **Aβ(1-42) tetramer** | | |
|---|---|---|---|---|---|---|
| | 0 | 1 | 2 | 0 | 1 | 2 |
| **+1** | 13543.1 | 15181.0 | 16818.9 | 18057.2 | 19695.0 | 21332.9 |
| **+2** | 6772.1 | 7591.0 | 8409.9 | 9029.1 | 9848.0 | 10667.0 |
| **+3** | 4515.0 | 5061.0 | 5607.0 | 6019.7 | 6565.7 | 7111.6 |
| **+4** | 3386.5 | 3796.0 | 4205.5 | 4515.0 | 4924.5 | 5334.0 |
| **+5** | 2709.4 | 3037.0 | 3364.6 | 3612.2 | 3939.8 | 4267.4 |
| **+6** | 2258.0 | 2531.0 | 2804.0 | 3010.4 | 3283.3 | 3556.3 |
| **+7** | 1935.6 | 2169.6 | 2403.6 | 2580.5 | 2814.4 | 3048.4 |
| **+8** | 1693.8 | 1898.5 | 2103.2 | 2258.0 | 2462.8 | 2667.5 |

***table S3:*** experimental values of the parameters tuned in the spectrometer to optimize the signal-to-noise ratio in MS experiments.

| Sample | Sampling Cone (V) | Trap (V) | Source Temp (°C) | Backing (mbar) |
|---|---|---|---|---|
| **Tetramer** | 50 | 50 | 40 | 5.85 |
| **Tetramer + SST14** | 50 | 50 | 40 | 5.85 |

***table S4:*** NMR titration points of the SST14 interaction study with βPFO$_{Aβ(1-42)}$ tetramer. For each experiment the concentration of SST14 is detailed as well as the corresponding rounded [SST14]:[Aβ(1-42) tetramer] ratio.

| | [SST14] (μM) | [SST14]:[βPFO$_{Aβ(1-42)}$ tetramer] |
|---|---|---|
| **control** | 0 | 0:1 |
| **T1** | 50 | 1:1 |
| **T2** | 99 | 2:1 |
| **T3** | 196 | 3:1 |
| **T3** | 291 | 5:1 |
| **T5** | 385 | 7:1 |
| **T6** | 566 | 10:1 |
| **T7** | 741 | 14:1 |
| **T8** | 909 | 17:1 |

#### Author Contributions

N.C. and A.R. designed and coordinated the project. E.P. prepared all the samples, acquired and analysed all the SEC, ITC, MS and NMR experiments. J.T. carried out and analysed the docking experiments. E.P. wrote the manuscript with input and contributions from all authors.
